# Intra-ocular Predation of Fluoroquinolone-Resistant *Pseudomonas aeruginosa* and *Serratia marcescens* by Predatory Bacteria

**DOI:** 10.1101/2023.09.17.558130

**Authors:** Eric G. Romanowski, Kimberly M. Brothers, Rachel C. Calvario, Nicholas A. Stella, Tami Kim, Mennat Elsayed, Daniel E. Kadouri, Robert M. Q. Shanks

## Abstract

Endogenous endophthalmitis caused by Gram-negative bacteria is an intra-ocular infection that can rapidly progress to irreversible loss of vision. While most endophthalmitis isolates are susceptible to antibiotic therapy, the emergence of resistant bacteria necessitates alternative approaches to combat intraocular bacterial proliferation. In this study the ability of predatory bacteria to limit intraocular growth of *Pseudomonas aeruginosa, Serratia marcescens*, and *Staphylococcus aureus* was evaluated in a New Zealand White rabbit endophthalmitis prevention model. Predatory bacteria *Bdellovibrio bacteriovorus* and *Micavibrio aeruginosavorus* were able to reduce proliferation of keratitis isolates of *P. aeruginosa* and *S. marcescens*. However, it was not able to significantly reduce *S. aureus*, which is not a productive prey for these predatory bacteria, suggesting that the inhibitory effect on *P. aeruginosa* requires active predation rather than an antimicrobial immune response. Similarly, UV-inactivated *B. bacteriovorus* were unable to prevent proliferation of *P. aeruginosa*. Together, these data suggest *in vivo* predation of Gram-negative bacteria within the intra-ocular environment.

## Introduction

Alternative approaches to antibiotics have become a major focus of research due to increasing resistance among bacterial pathogens. One avenue of research for this purpose is the use of predatory bacteria as a “living antibiotic” [1-4]. Predatory bacteria such as *Bdellovibrio bacteriovorus* and *Micavibrio aeruginosavorus* use a wide range of Gram-negative bacteria as a food source including numerous pathogens [5]. It has been demonstrated that predatory bacteria are indifferent to the antibiotic-resistance status of their prey [6, 7]. Moreover, it has been demonstrated that predatory bacteria are non-toxic to mammalian cells and in animal models and have the ability to attenuate Gram-negative bacteria several *in vivo* models, including but not limited to airway and oral infections in rodents and central nervous system infection of zebrafish hindbrain ventricles [8-16].

Recent studies have evaluated the use of predatory bacteria for treatment of ocular surface infections [17-19]. These have demonstrated some efficacy against *Escherichia coli* [17] in a mouse model and *Pseudomonas aeruginosa* in rabbit models including the prevention of corneal perforations [19]. However freeze-dried *B. bacteriovorus* proved to be ineffective in treating *Moraxella bovis* in a large animal keratoconjunctivitis model [20]. The prior studies have focused on the anterior portion of the eye. In this study the goal was to evaluate the use of predatory bacteria in preventing bacterial proliferation in an endophthalmitis model using New Zealand White rabbits.

Endogenous endophthalmitis is frequently caused by Gram-negative bacteria that travel from the blood stream, through the blood-brain barrier and into the posterior portion of the eye [21, 22]. There bacteria can rapidly proliferate, induce a damaging immune response, and cause damage to the tissues crucial for vision such as the retina. This can lead to severe surgical interventions such as removal of the eyes content (evisceration) or the entire globe (enucleation). Endophthalmitis caused by Gram-negative bacteria such as *Klebsiella pneumonia* is especially prominent in Asia [21-23]. At our hospital the most frequent Gram-negative bacteria that cause endophthalmitis are *Pseudomonas aeruginosa* and *Serratia marcescens*, whereas bacteria from the *Staphylococcus* and *Streptococcus* genera are the most prominent overall [24].

In this study, the ability of *B. bacteriovorus* strain HD100 and *M. aeruginosavorus* strain ARL-13 were evaluated for the ability to prevent growth of bacterial pathogens within the eye and reduced proliferation of *P. aeruginosa* was reproducibly observed. Furthermore, whether inhibition of *P. aeruginosa* was due to active predation or an immune response triggered by predatory bacteria, was tested using ultra violet light inactivated *B. bacteriovorus*. Data suggests that both tested predatory bacteria prey upon *P. aeruginosa* within the vitreous humor of the rabbit eye.

## Materials and Methods

### Strains and bacterial growth conditions

The predatory bacteria *Bdellovibrio bacteriovorus* HD100 (ATCC 15356) [25, 26] and *Micavibrio aeruginosavorus* ARL-13 [27] were used in the study. Predator lysates (co-cultures) were prepared as previously reported [28]. Predators *B. bacteriovorus* and *M. aeruginosavorus* were incubated at 30°C with *E. coli* strain WM3064 [29] (1 × 10^9^ CFU/ml) for 24 and 72 hours respectively. The resulting lysates were passed twice through a 0.45-μm pore-size filter (Millipore, Billerica, MA, USA). Predators were washed with phosphate buffered saline (PBS) and concentrated by three sequential 45-minute centrifugations at 29,000 x g. Finally, predator pellets were suspended in PBS to reach a final concentration of 1.7 × 10^10^ PFU/ml *B. bacteriovorus* and 3.5 × 10^9^ PFU/ml *M. aeruginosavorus. B. bacteriovorus* UV inactivated cells were prepared by placing 1 ml of purified predator sample in a well of a 12 well plate and radiating the plate 20 times on the Auto Cross Link setting, while mixing the sample in-between each cross-link (UV Stratalinker 1800; Stratagene, San Diego, CA, USA). Lack of predator viability was confirmed by PFU plating, in which no plaque had developed. Structural integrity of the predator cells was confirmed by light microscopy (1000x magnification).

Pathogens used for this study were *P. aeruginosa* strain PaC, which is a fluoroquinolone resistant ocular isolate [30]. *S. marcescens* strain K904 is a keratitis ocular isolate [31]. *S. aureus* strain E277 is an endophthalmitis isolate from The Charles T. Campbell Laboratory deidentified strain collection. These were streaked to single colonies on TSA blood agar plates from stocks stored at −80°C. Single colonies were grown with aeration in lysogeny broth for 16-18h and then adjusted in PBS to an inoculum of 5-10 x 10^3^ CFU in 25 μl for injection into eyes.

### Animal experiments

This study was approved by the University of Pittsburgh’s Institutional Animal Care and Use Committee (Protocol #15025331 “The Use of Predatory Bacteria to Treat Ocular Infections”) and conformed to the ARVO Statement on the Use of Animals in Ophthalmic and Vision Research.

Female New Zealand White rabbits weighing 1.5-2 kg were received from Charles River Laboratories’ Oakwood Rabbitry. Following systemic anesthesia with 40 mg/kg of ketamine & 4 mg/kg of xylazine administered intramuscularly, and topical anesthesia with 0.5% proparacaine, the right eyes were inoculated in the vitreous via pars plana injection with 25 μl of the bacterial suspension or PBS depending on the group. Immediately following injections of the bacteria, the same eyes were injected with 0.1 ml (100 μl) of the predatory bacteria or PBS per the experimental groups above. Injection of the predatory bacteria or PBS was performed in a different location than the pars plana injection of bacteria. Rabbits were treated with 1.5 mg/kg ketoprofen, administered intramuscularly, to reduce pain after recovery from anesthesia. At 24 hours after inoculation, the rabbits were examined using a slit lamp and imaged. Rabbits were euthanized with an overdose of intravenous Euthasol solution following systemic anesthesia with ketamine and xylazine administered intramuscularly. Vitreous humor taps were performed on the infected eyes by inserting a 23-gauge needle attached to a 1 cc syringe into the vitreous chamber about 4 mm from the limbus and removing about 0.2-0.3 ml of fluid. The vitreous humor was transferred to a sterile tube and placed on ice. Standard colony counts determinations were performed on the vitreous samples using 5% sheep blood agar plates and incubated overnight at 37°C. Cytokines were detected from vitreous humor using commercial kits IL-1β (Sigma-Aldrich), TNFα (Thermo Scientific). Clinical signs of endophthalmitis were determined by a masked reviewer and used a 10-point scale with a 0-2-point score given for discharge, redness of eye, chemosis, anterior eye involvement, and hypopyon.

### Statistical analysis

Data was analyzed using GraphPad Prism software. All experiments were repeated at least twice. Kruskal-Wallis with Dunn’s post-test was used to compare medians and ANOVA with Tukey’s post-test was used to analyze means.

## Results

### Predatory bacteria prevent proliferation of *P. aeruginosa*

*P. aeruginosa* strain PaC is a fluoroquinolone resistant ocular clinical isolate that was chosen because it is susceptible to predation by predatory bacteria [32]. The ocular vitreous chamber of NZW rabbits were injected with *P. aeruginosa* (5.0 x 10^3^ CFU) followed by *B. bacteriovorus* (4.3 x 10^8^ PFU) or *M. aeruginosavorus* (8.8 x 10^7^ PFU). Controls included injection of vehicle (PBS) or individual microbes. At 24 hours eyes were examined by a slit lamp, imaged, and graded for clinical signs of inflammation. Eyes injected with predatory bacteria had increased inflammatory scores that were not significantly higher than the vehicle only control (PBS + PBS), Figure 1A. Eyes injected with *P. aeruginosa* only had a notable increase in inflammatory score (p<0.0001), by comparison, ocular inflammation was reduced in eyes injected with both *P. aeruginosa* and either predatory bacteria. Although, the clinical signs of inflammation were not significantly different between the PaC alone and the PaC with predatory bacteria, the eyes injected with both PaC and *B. bacteriovorus* HD100 were not significantly worse than the vehicle control eyes (Figure 1A). Representative images of eyes from each group are shown in Figure 1B.

**Figure 1.**
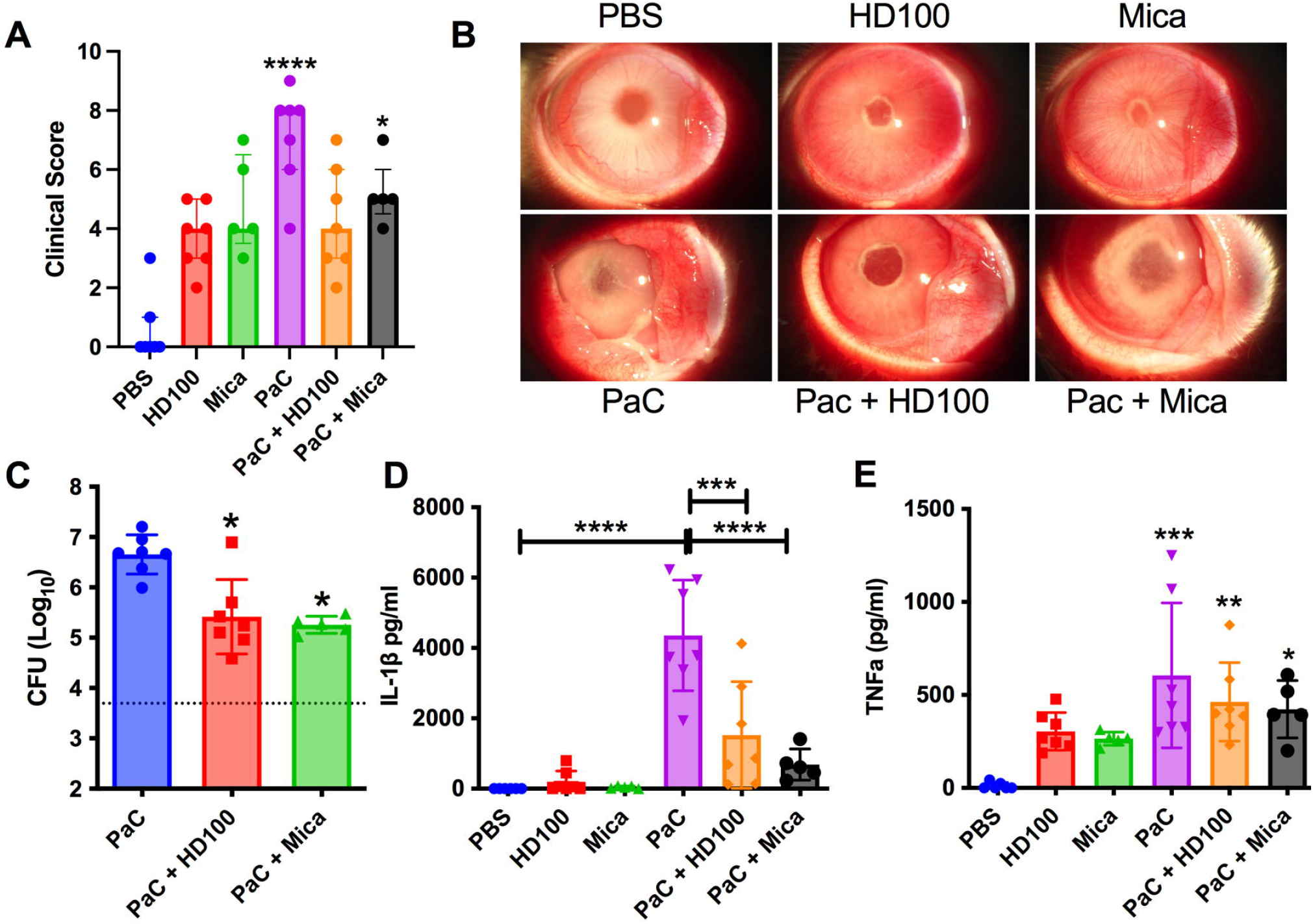
Effect of predatory bacteria on inflammation and *P. aeruginosa* (PaC) proliferation in a rabbit model of endophthalmitis. Each point represents a rabbit in all figures. HD100 indicates *B. bacteriovorus*; Mica signifies *M. aeruginovorus*. **A**. Clinical inflammation scores for rabbits 24 hours post-inoculation. Only the *P. aeruginosa* group (PAC) and *P. aeruginosa* with *M. aeruginovorus* groups were statistical different from the vehicle control, n=5-7. Medians and interquartile ranges (IQR) are shown. Asterisks indicate statistical differences from the PBS group. **B**. Representative eyes from subject with median clinical scores are shown for each group. **C**. Intraocular CFU/ml at 24h post-injection (Median and IQR, n=5-7). Asterisks indicate statistical differences from the PaC group, *, p<0.05. The dotted line indicates the inoculum of PaC introduced into the eye. **D**. IL-1β cytokine measured by ELISA, means and standard deviations (SD) are shown, (n=5-7). **E**. TNFα cytokine measured by ELISA, means and standard deviations (SD) are shown, (n=5-7). Asterisks indicate differences from the PBS group. *, p<0.05; **, p<0.01; ***, p<0.001; ****, p<0.0001.

The trend toward reduced inflammation following addition of predatory bacteria to *P. aeruginosa* treated eyes correlated with a >95% reduction in the *P. aeruginosa* bacterial burden in eyes injected with both predatory bacteria and *P. aeruginosa* compared to *P. aeruginosa* alone, p<0.05 (Figure 1C). Unchecked, *P. aeruginosa* replicated from an inoculum of 5 x 10^3^ to a burden 4.8 x 10^6^ CFU in 24h in the eye. This was 17-fold and 25-fold higher than what it achieved with *B. bacteriovorus* HD100 and *M. aeruginosavorus* respectively.

Damage associated pro-inflammatory cytokine IL-1β levels followed a similar trend to the clinical scores and were largely unaffected by predatory bacteria alone, elevated with *P. aeruginosa* alone, and significantly mitigated in eyes with both predatory bacteria and *P. aeruginosa* (Figure 1D). A matching trend for pro-inflammatory cytokine TNFα was measured, although to a lesser extent where the predatory bacteria did not significantly reduce the cytokine levels compared to the *P. aeruginosa* alone (Figure 1E).

### Predatory bacteria reduce intraocular proliferation of *S. marcescens*

Figure 2 shows experimental data in which predatory bacteria were evaluated for their ability to prevent intraocular growth of *S. marcescens*. Clinical presentation of *S. marcescens* infected eyes was less severe than that of *P. aeruginosa* and not notably changed by the addition of predatory bacteria (Fig 2A). Representative images are shown in Figure 2B. *S. marcescens* was able to grow 85,800-fold from 5.6 x 10^3^ inoculum to 4.8 x 10^8^ intraocularly in 24 hours. This was reduced 2.2 and 2.7-fold by *B. bacteriovorus* and *M. aeruginosavorus* respectively, and was significantly reduced by *B. bacteriovorus* (Fig 2C).

**Figure 2.**
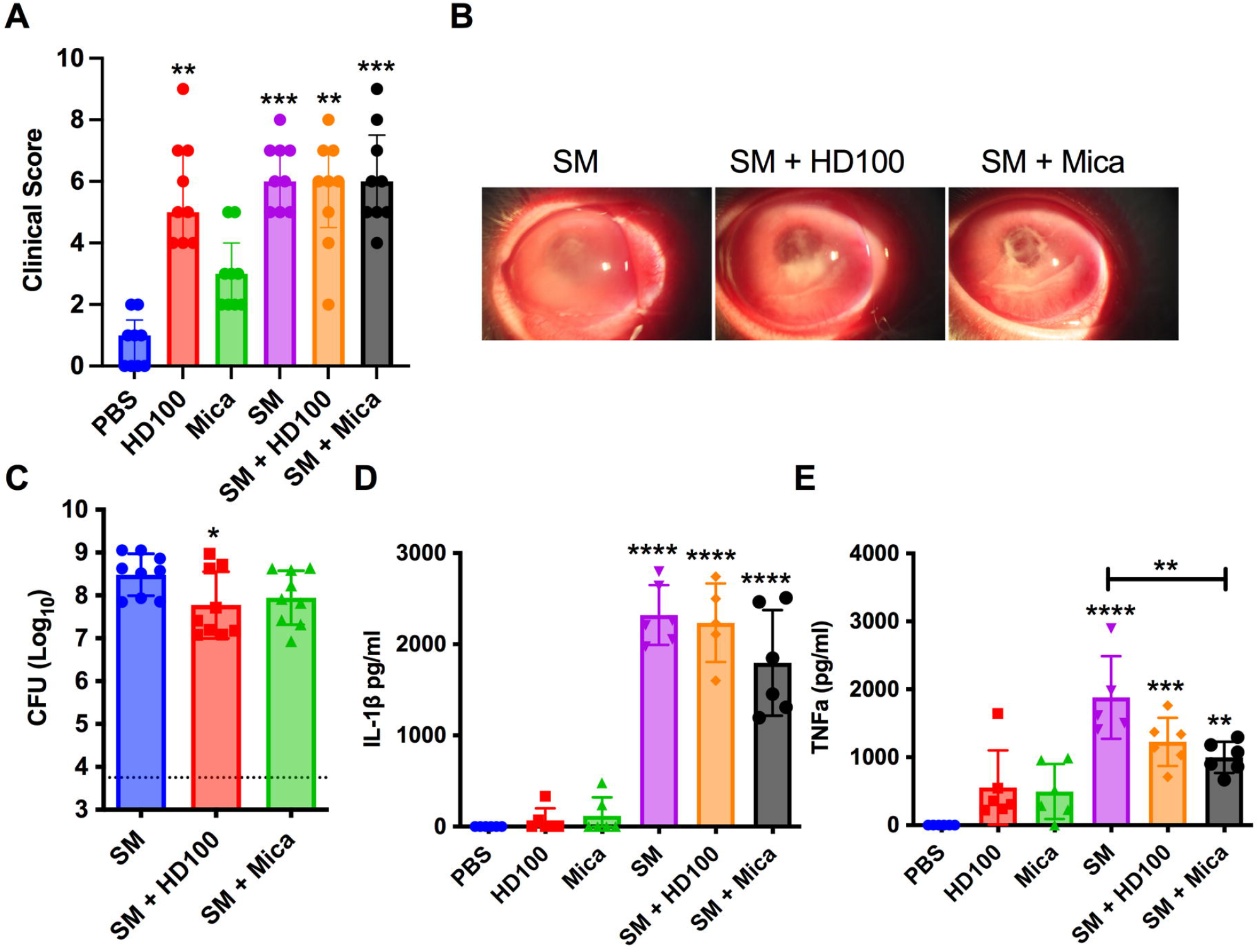
Impact of predatory bacteria on *S. marcescens* (SM) proliferation and consequent cytokine production. **A**. Clinical inflammation scores for rabbits 24 hours post-inoculation. n=9. Medians and interquartile ranges (IQR) are shown. Asterisks indicate statistical differences from the PBS group. **B**. Representative images with median clinical scores. **C**. Intraocular CFU/ml at 24h post-injection (Median and IQR, n=9). Asterisks indicate statistical differences from the SM group (p<0.05). The dotted line indicates the inoculum of *S. marcescens* injected into the eye. **D**. IL-1β cytokine measured by ELISA, means and standard deviations (SD) are shown, (n=5-8). **E**. TNFα cytokine measured by ELISA, means and standard deviations (SD) are shown, (n=5-8). Asterisks indicate differences from the PBS group, or where indicated by brackets. *, p<0.05; **, p<0.01; ***, p<0.001; ****, p<0.0001.

Similar to clinical presentation, IL-1β levels were induced less by *S. marcescens* than *P. aeruginosa* and were unaffected by co-incubation with predatory bacteria (Figure 2D). By contrast, TNFα levels were higher in *S. marcescens* infected eyes than *P. aeruginosa* and were significantly reduced when *S. marcescens* was co-incubated with *M. aeruginosavorus* (Figure 2E).

### Predatory bacteria do not prevent *S. aureus* replication in the eye

Although *B. bacteriovorus* was shown to attach to the Gram-positive bacterium *S. aureus, S. aureus* does not support *B. bacteriovorus* or *M. aeruginosavorus* full predation or growth cycle [25, 33-35]. Inclusion of this bacteria should therefore give insight into whether the prevention of *P. aeruginosa* and *S. marcescens* intraocular growth by predatory bacteria is due to active predation or other mechanism, such as predatory bacteria-induced biosynthesis of antimicrobials or immune response triggered by the presence of the predators. The clinical evaluation showed modest inflammation with *S. aureus* alone that was not significantly altered with addition of predatory bacteria (Fig 3A). Unchallenged by predatory bacteria, the *S. aureus* strain was able to grow 87-fold from 1.0 x 10^4^ to 8.9 x 10^5^ in 24 hours. *B. bacteriovorus* was associated with a 2.26-fold reduction in *S. aureus* burden, whereas *M. aeruginosavorus* reduced less than 2-fold (19%) and neither of these changes were significantly different by ANOVA (Figure 3B). IL-1β levels were not reduced by the presence of predatory bacteria (Figure 3C).

**Figure 3.**
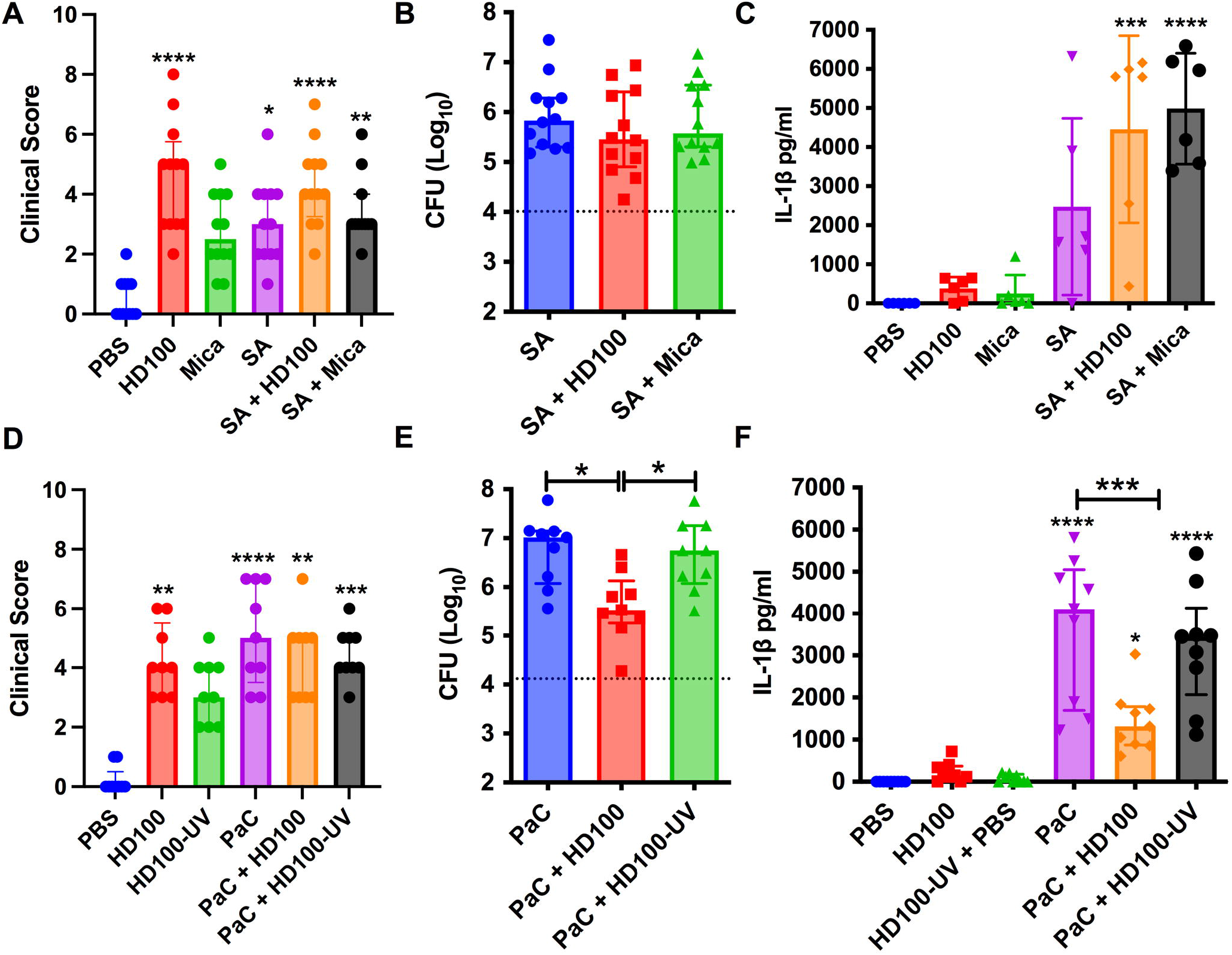
Differentiation between predation and induced antimicrobial host-response. **A, D**. Clinical inflammation scores for rabbits 24 hours post-inoculation. n=9-12. Medians and interquartile ranges (IQR) are shown. Asterisks indicate statistical differences from the PBS group. SA indicates *S. aureus* **B, E**. Intraocular CFU/ml at 24h post-injection (Median and IQR, n=9-12). Asterisks indicate statistical differences from the SA and PaC groups. The dotted line indicates the initial inoculum. **C, F**. IL-1β cytokine measured by ELISA, means and standard deviations (SD) are shown, (n≥6). Asterisks indicate differences from the PBS group, or where indicated by brackets. *, p<0.05; **, p<0.01; ***, p<0.001; ****, p<0.0001.

### Viable predatory bacteria are required to reduce intraocular proliferation of *P. aeruginosa*

To further test whether the reduction in *P. aeruginosa* was mediated by predation, UV-inactivated *B. bacteriovorus* were tested. The UV-inactivated bacteria correlated with modestly increased clinical scores (Figure 3D). Importantly, the *P. aeruginosa* inhibition by live *B. bacteriovorus* was absent for eyes using the UV-inactivated predator (Figure 3E). Similarly, the reduction in *P. aeruginosa* induced IL-1β was reduced by live but not inactivated predatory bacteria (Figure 3F).

## Discussion

This proof-of-principle study determined whether predatory bacteria could prey upon bacteria within the eye. This study demonstrated a clear ability of both tested predatory bacteria to reduce *P. aeruginosa* proliferation within the eye. While the reduction was less with *S. marcescens*, predation may be masked by the remarkably fast replication of *S. marcescens* strain K904 in the eye. While two different experiments differed in the extent to which *P. aeruginosa* caused clinical inflammatory signs (Figure 1A versus Figure 3A), the IL-1β levels were reproducibly reduced when predatory bacteria were added to *P. aeruginosa* infected eyes. IL-1 is associated with ocular tissue damage that triggers production of pro-inflammatory cytokines and reduces barrier function [36]. IL-1β is a marker of bacterial endophthalmitis associated with productive antimicrobial host-responses [37-39]. Unlike the tested pathogens, predatory bacteria failed to significantly induce intraocular cytokine levels above that of eyes injected with the PBS vehicle. This is in agreement with prior *in vitro* and *in vivo* studies [12, 13, 19, 20, 28, 32, 40-42] and, at least for *B. bacteriovorus* due to its unusual outer membrane composition and membrane-sheathed flagellum which reduce TLR4 and TLR5 activation [43, 44], which are major mediators of inflammation in bacterial endophthalmitis [45, 46].

Data from this study support the hypothesis that predatory bacteria actively prey upon *P. aeruginosa* and possibly *S. marcescens* in the vitreous chamber of the eye. This is based on two observations. First is that predatory bacteria failed to significantly reduce the proliferation of a bacteria that they are unable to prey upon, which suggests that an antimicrobial host response is not strongly induced by predatory bacteria. The second being that UV-inactivated predatory bacteria were unable to reduce intraocular *P. aeruginosa*. While innate ocular defense mechanisms such as neutrophils likely play a role in inhibition of pathogen growth, the reduced proliferation of *P. aeruginosa* and *S. marcescens* in vitreous chamber of the eye in the eyes co-incubated with predatory bacteria is most likely due to active predation. Although our findings do support the hypothesis that active predation is required for the effect seen in Gram-negative pathogens, one cannot rule out that an elevated immune response might be triggered following predation as prey cell debris accumulates. A synergistic effect between the immune system and *Bdellovibrio* was previously suggested in a study monitoring predation of *Shigella* using a Zebrafish infection model [16].

In conclusion, this study strongly suggests that predatory bacteria can kill bacteria in the vitreous chamber of the eye. However, on its own would likely be insufficient to treat a clinical endophthalmitis. Further studies to determine whether predatory bacteria coupled with antibiotics would improve antimicrobial outcomes as has been previously tested *in vitro* [47].

## Acknowledgements

This study was funded by National Institutes of Health grants R01*EY027331* (to R.S.) and CORE Grant P30 EY08098 to the Department of Ophthalmology. The Eye and Ear Foundation of Pittsburgh and from an unrestricted grant from Research to Prevent Blindness, New York, NY provided additional departmental funding. This work was also funded by the U.S. Army Research Office and the Defense Advanced Research Projects Agency (DARPA) and was accomplished under Cooperative Agreement Number W911NF-15-2-0036 to DEK and RMQS. The views and conclusions contained in this document are those of the authors and should not be interpreted as representing the official policies, either expressed or implied, of the Army Research Office, DARPA, or the U.S. Government. The U.S. Government is authorized to reproduce and distribute reprints for Government purposes notwithstanding any copyright notation hereon.

